# SNP- Based Assessment of Genetic Purity and Diversity in Maize Hybrid Breeding

**DOI:** 10.1101/2021.03.22.436406

**Authors:** Chimwemwe Josia, Kingstone Mashingaidze, Assefa B. Amelework, Aleck Kondwakwenda, Cousin Musvosvi, Julia Sibiya

## Abstract

Assessment of genetic purity of inbred lines and their resultant F_1_ hybrids is an essential quality control check in maize hybrid breeding, variety release and seed production. In this study, genetic purity, parent-offspring relationship and diversity among the inbred lines were assessed using 92 single-nucleotide polymorphism (SNP) markers. A total of 188 maize genotypes, comprising of 26 inbred lines, four doubled haploid (DH) lines and 158 single-cross maize hybrids were investigated in this study using Kompetitive Allele Specific Polymerase Chain Reaction (KASP) genotyping assays. The bi-allelic data was analyzed for genetic purity and diversity parameters using GenAlex software. The SNP markers were highly polymorphic and 90% had polymorphic information content (PIC) values of > 0.3. Pairwise genetic distances among the lines ranged from 0.05 to 0.56, indicating a high level of dissimilarity among the inbred lines. Maximum genetic distance of (0.56) was observed for CKDHL0089, CML443 and CB323, while the lowest (0.05) was between I-42 and I-40. The majority (67%) of the inbred lines studied were genetically pure with residual heterozygosity of <5%, while only 33% were had heterozygosity levels of >5%. Inbred lines, which were not pure, require purification through further inbreeding. Cluster analysis partitioned the lines into three distinct genetic clusters with the potential to contribute new beneficial alleles to the maize breeding program. Out of the 68 hybrids (43%) that passed the parent-offspring test, seven hybrids namely; SCHP29, SCHP95, SCHP94, SCHP134, SCHP44, SCHP114 and SCHP126, were selected as potential candidates for further evaluation and release due to their outstanding yield performance.

## 1.1 Introduction

Maize (*Zea mays* L.) is the principal crop for food security and nutrition for the majority of people in sub-Saharan Africa (SSA) and Latin America [1]. However, adequate production of the crop is hampered by low grain yields due to biotic and abiotic stresses. Hence, there is a need to improve grain yield through hybrid breeding to exploit heterosis.

Assessment of genetic purity of parental inbred lines and parent-offspring test for the resultant F_1_ hybrids is an essential quality control function in maize hybrid breeding programs. These functions are now more critical due to the stringent intellectual property requirements governing plant breeding and variety registration in many countries [2]. Additionally, the maintenance of high levels of genetic purity is critical for the robust agronomic performance of the genotype. Parent-offspring test helps to prove parentage for a specific hybrid whether it is a true derivative of the original parental inbred lines without pollen contamination [3]. Inbred lines’ genetic purity and parentage can be proved using three approaches namely; grow out test (GOT), use of biochemical markers and use of molecular markers.

Grow out test is a morphologically based approach using a set of descriptors, while the biochemical markers approach analyses the protein/isoenzyme profiles of the genotype and the molecular marker approach detects variation of the genotype directly at the DNA level [3]. Unlike GOT and biochemical marker methods, which have low polymorphism and high environmental influences, molecular markers are ideal for genotyping since they are codominant, highly abundant and polymorphic, independent of the environment, reproducible, expressed at all developmental stages, known position in the genome, linked to traits of interest and automation is possible [3].

Semagn et al. [4] highlighted several types of molecular markers that are available for detection of polymorphism. The main ones include; restriction fragment length polymorphism (RFLPs), random amplified polymorphic DNA (RAPD), simple sequence repeats (SSRs), amplified fragment length polymorphism (AFLP) and single nucleotide polymorphism (SNP). In this study, SNP markers were used to determine the genetic purity of the maize parental inbred lines and to prove the parentage of the resultant single-cross hybrids. Recent advances in molecular technology have emphasized the use of SNP markers because they are cost-effective per data point, adequate genomic abundance, locus-specificity, codominant, and potential for high throughput unlike the other markers [5].

The application of molecular markers is more efficient, saves time and resources [6] and they are free from environmental influences compared to morphological markers. It is often assumed that the use of a large number of markers results in higher accuracy. In most sequence based marker systems, the levels of missing data can lead to wrong interpretation, hence, the selection of fewer markers with high and repeatable representation across samples is desired and is cost-effective. Chan et al. [7] suggested that fewer markers with high excepted heterozygosity, missing value of <20%, and observed heterozygosity of <6% are ideal markers for accurate quality control genotyping. Similarly, Semagn et al [2] suggested that a set of 50-100 single plex assay SNPs are adequate for molecular-based quality control genotyping. It is against this background that in the present study, a set of 92 SNPs were used to genotype 30 parental lines and 158 single-cross hybrids.

According to Gowda et al. [3], parental inbred lines are expected to be pure with residual heterozygosity of less than 5%. Inbred lines having residual heterozygosity above 5% are either not pure due to genetic contamination or not fixed unless if they were deliberately maintained at early generation during development. Genetic contamination reduces the genetic and physiological quality of the seeds leading to decreased crop productivity [8]. Hence, inbred line genetic purity assessment and parent offspring test are important quality control procedures for a successful hybrid breeding program.

## 1.2 Materials and methods

### 1.2.1 Experimental material

A total of 188 maize genotypes, comprising 26 elite parental inbred lines, four doubled haploid lines and 158 experimental single-cross hybrids (Supplementary Table 1) were genotyped using 92 single nucleotide polymorphism (SNP) markers (Supplementary Table 2). These markers are a subset of the 100 SNP markers recommended by CIMMYT for routine quality control genotyping in maize and cover the 10 pairs of the maize chromosomes [3]. All the genotypes used in this study were sourced from Agriculture Research Council-Grain Crops Institutes (ARC-GCI), Potchefstroom, South Africa.

### 1.2.2 Leaf sampling, hybrids’ field evaluation, DNA extraction and genotyping

Maize genotypes were planted at ARC-GCI research farm, Potchefstroom (26°74”S; 27°8’E) during the 2017/18 summer season. Leaf sampling was done using the supplied LGC sampling kit (LGC Genomics Laboratory, United Kingdom). Five to eight leaf discs were taken per entry five weeks after planting for DNA extraction. Leaf samples from the same entry were placed in a specific 2 x 96-well plate with each well representing an individual genotype. Each well was sealed using a perforated trip cap and the desiccant sachet was placed directly on top of the strip cap-sealed well and the plastic lid was replaced on top. The storage rack was secured using an elastic band and was placed inside a sealable plastic bag. The sealed bag was placed into the plant kit box and the samples were shipped to LGC Genomics Laboratory, in the United Kingdom for genotyping. DNA extraction, amplification and visualization were done according to the LGC protocol (www.lgcgroup.com). Genomic DNA was extracted from the leaf disc samples and the quality and quantity of the extracted DNA was determined before genotyping. Genotyping was done using the 92 SNP markers, following the Kompetitive Allele Specific Polymerase Chain Reaction (KASP) protocol used by LGC Genomics (www.lgcgroup.com).

Field evaluation of single-cross hybrids was done at three locations namely; Potchefstroom (ARC-GCI) in North West province, Cedara in KwaZulu-Natal province and Vaalharts in the Northern Cape province, South Africa, during the 2017/18 summer season. The trial constituted of five production environment; Potchefstroom and Cedara representing two environments (low N and optimum), while Vaalharts had an optimum environment. One hundred and seventy single-cross maize hybrids including three checks were laid out in a 34 x 5 (0,1) alpha lattice experimental design replicated twice in each environment. However, for the purpose of this study only 158 set of single cross hybrids were used due to limited availability of seeds for the dropped hybrids.

### 1.2.3 Statistical analyses

Data filtering for monomorphic SNPs and/or SNPs with a minor allele frequency of less than 2% were performed and all the 92 SNPs were polymorphic and of high quality. Genetic purity of the parental inbred lines was calculated as percentage residual heterozygosity using the formula shown in equation (1) described by Gowda et al. [3]. Genetic diversity parameters such as observed heterozygosity (Ho), expected heterozygosity (He), and fixation index (F_IS_) were determined using GenAlex version 6 [9]. The formula: PIC =1 − ΣP_ij_^2^, where P_ij_ is the frequency of j^th^ allele of the i^th^ locus, were used to calculate Polymorphic information content (PIC).

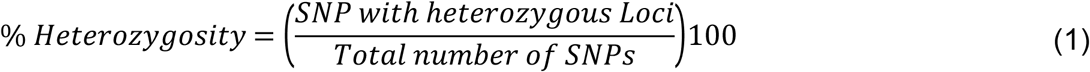

Genetic relationships within and among the inbred lines were assayed with a neighbor-joining algorithm, using the unweighted pair group method (UWPGM) in DARwin 6.0 software [10]. Pairwise dissimilarity matrices were obtained from the Jaccarrd’s coefficient and a dendrogram was generated. For the node construction, a bootstrap analysis was performed, based on 10000 bootstrap values, using DARwin. The distinctiveness of the clusters was checked, using the cophenetic correlation coefficient (r). The parent-offspring relationship for each parent-hybrid pair was tested according to methods described by Gowda et al. [3]. Parameters such as the proportion of SNPs from parent A and parent B, SNPs shared by both parents and SNPs that do not belong to either of the parent were estimated.

## 1.3 Results

### 1.3.1 SNP characterization

The distributions of values for polymorphic information content, gene diversity, inbreeding coefficient and minor allele frequencies of the 92 SNPs estimated on the 188 maize genotypes are shown in Figure 1. Inbreeding coefficient displayed contrasting values ranging from −0.17 to 1.00, with a mean of 0.10. About 24% of the SNPs showed negative F_IS_ values. Nearly 39% of the SNPs had F_IS_ values between 0.10 and 0.40 (Figure 1a). The SNPs diversity ranged from 0.11 to 0.50; however, the vast majority (92%) fell between 0.30 and 0.50 and eight SNPs revealed moderate gene diversity (Figure 1b). Approximately 90% of the markers used in this study had PIC values exceeding 0.30. The majority of the values (77%) were between 0.40 and 0.50 and only one marker (PZA03527_3) displayed a PIC value of less than 0.2 (Figure 1c). The minor allele frequency ranged from 0.06 for the marker PZA03527_3 to 0.50 for the marker sh1_12, with a mean of 0.35 (Figure 1d). More than 55% of the SNPs revealed a minor allele frequency exceeding the mean (0.35). Observed heterozygosity (H_o_) values ranged from 0.0 to 0.56 with a mean of 0.40 (data now shown). SNP markers PZA00793_2 and PHM2350_17 had H_o_ value of 0.0 indicating the alleles of these SNPs were 100% fixed among the maize genotypes, however, 97% of the SNPs had H_o_ values exceeding 15%. SNP markers PHM2350_17, and PZA00793_2 showed inbreeding coefficient value of 1.

**Figure 1.**
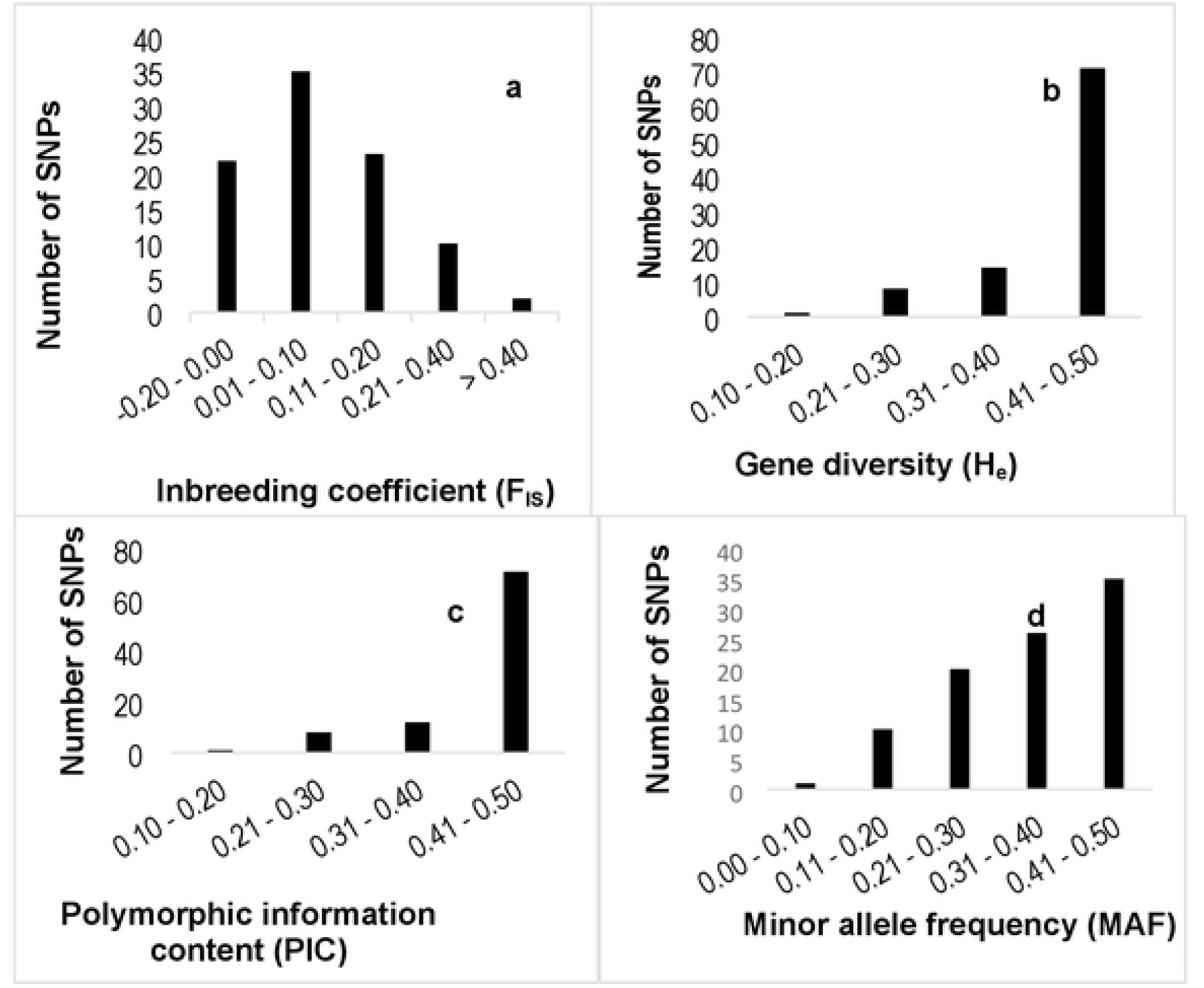
Distribution of the 92 SNPs estimated for all 188 maize genotypes for (a) Inbreeding coefficient, (b) Gene diversity, (c) Polymorphic information content and (d) Minor allele frequency

The genetic diversity parameter estimates of the 92 SNPs used in this study summarized per chromosome are presented in Table 1. The number of SNPs on each chromosome ranged from six on chromosome 10 to 12 on chromosomes 2 and 5, with a mean of 9.2 SNPs per chromosomes. The observed heterozygosity of the SNP loci for the inbred lines ranged from 6% to 11%, while the hybrids revealed H_0_ values ranging from 39% to 49%. The gene diversity values for the inbred lines ranged from 0.42 to 0.49, with a mean gene diversity of 0.45. However, no significant differences were observed in PIC and gene diversity values among the ten chromosomes. The mean inbreeding coefficient (F_IS_) was significantly higher for inbred lines ranging from 0.75 to 0.85, with a mean value of 0.80. The hybrids on the other hand, revealed very low F_IS_ values ranging from −0.02 to 0.17, with a mean value of −0.05.

**Table 1:** Summary of genetic diversity parameters of 92 SNPs per chromosome measured in a set of 188 maize genotypes

| Chromosome | No. SNPs | Genetic parameters |  |  |  |  |  |  |  |
| --- | --- | --- | --- | --- | --- | --- | --- | --- | --- |
|  |  | Maize Inbred lines |  |  |  | Hybrids |  |  |  |
|  |  | H<sub>o</sub> | H<sub>e</sub> | F<sub>IS</sub> | PIC | H<sub>o</sub> | H<sub>e</sub> | F<sub>IS</sub> | PIC |
| 1 | 9 | 0.07 | 0.42 | 0.82 | 0.42 | 0.41 | 0.40 | -0.03 | 0.40 |
| 2 | 12 | 0.11 | 0.45 | 0.75 | 0.44 | 0.44 | 0.42 | -0.05 | 0.42 |
| 3 | 11 | 0.08 | 0.45 | 0.80 | 0.44 | 0.47 | 0.44 | -0.07 | 0.43 |
| 4 | 9 | 0.07 | 0.45 | 0.84 | 0.44 | 0.45 | 0.45 | -0.02 | 0.45 |
| 5 | 12 | 0.11 | 0.46 | 0.76 | 0.45 | 0.51 | 0.45 | -0.13 | 0.45 |
| 6 | 9 | 0.09 | 0.45 | 0.79 | 0.44 | 0.44 | 0.42 | -0.05 | 0.42 |
| 7 | 7 | 0.10 | 0.42 | 0.75 | 0.30 | 0.49 | 0.44 | -0.12 | 0.44 |
| 8 | 9 | 0.08 | 0.49 | 0.84 | 0.49 | 0.39 | 0.43 | 0.17 | 0.43 |
| 9 | 8 | 0.06 | 0.44 | 0.85 | 0.43 | 0.46 | 0.42 | -0.08 | 0.42 |
| 10 | 6 | 0.09 | 0.46 | 0.81 | 0.45 | 0.47 | 0.44 | -0.06 | 0.44 |
| Overall mean |  | 0.09 | 0.45 | 0.80 | 0.44 | 0.45 | 0.43 | -0.05 | 0.43 |
| SE |  | 0.00 | 0.01 | 0.01 | 0.01 | 0.01 | 0.01 | 0.02 | 0.01 |

N= Number of individuals tested; H<sub>o</sub>= observed heterozygosity; H<sub>e</sub>= expected heterozygosity;
F<sub>IS</sub>= inbreeding coefficient; PIC= polymorphic information content; SE= Standard error

### 1.3.2 Genetic purity of parental maize inbred lines

The percentage of missing data per SNP in this study was below 3% and varied from 0 to 2.17%, with the overall mean of 1.16%. Based on the 92 SNPs, genetic purity among the 30 inbred lines varied from 0.0 to 57.6, with an overall mean of 10.43 (Table 2). All the 92 SNP loci tested in this study were fixed in 60%. Out of the 18 genotypes that showed 100% genetic purity, four lines (CKDHL0089, CKDHL0295, CKDHL0378 and CKDHL0470) were doubled haploids. Inbred lines CKL05022 and CB323 had a heterozygous percentage of less than 5% and these inbred lines were considered to be fixed. However, 33.3% of the inbred lines had residual heterozygosity ranging from 5.43 to 57.61%.

**Table 2:** Genetic purity of 26 maize inbred lines and four doubled haploid lines based on 92 SNPs

| Name | % of missing alleles | % of heterozygote alleles |
| --- | --- | --- |
| CKDHL0089 | 2.17 | 0.00 |
| CKDHL0295 | 0.00 | 0.00 |
| CKDHL0378 | 1.09 | 0.00 |
| CKDHL0470 | 0.00 | 0.00 |
| CML202 | 1.09 | 0.00 |
| CML216 | 1.09 | 0.00 |
| CML442 | 1.09 | 0.00 |
| CML443 | 1.09 | 0.00 |
| CB322 | 2.17 | 0.00 |
| CML544 | 0.00 | 0.00 |
| CML511 | 0.00 | 0.00 |
| CZL068 | 0.00 | 0.00 |
| I-38 | 2.17 | 0.00 |
| I-42 | 2.17 | 0.00 |
| CML540 | 2.17 | 0.00 |
| CZL99017 | 1.09 | 0.00 |
| CML312 | 2.17 | 0.00 |
| I-40 | 1.09 | 0.00 |
| CKL05022 | 1.09 | 2.00 |
| CB323 | 2.17 | 4.35 |
| CK21 | 1.09 | 5.43 |
| RO549W | 2.17 | 15.22 |
| U2540W | 1.09 | 16.30 |
| CML444 | 2.17 | 18.48 |
| CB339 | 0.00 | 19.57 |
| CML543 | 2.17 | 39.13 |
| CZL0919 | 1.09 | 42.39 |
| CML488 | 0.00 | 45.65 |
| CML547 | 0.00 | 48.91 |
| CZL0718 | 1.09 | 57.61 |
| Mean | 1.16 | 10.43 |
| SE | 0.16 | 3.23 |

### 1.3.3 Genetic relationship among 30 parental lines

The population structure for the parental lines was assessed using distance-based cluster analyses. Cluster analysis based on Jaccard’s genetic distance values classified the 30 parental lines into three distinct clusters (Figure 2). The distinctiveness of the clusters was confirmed by the high cophenetic correlation coefficient for SNPs (r = 0.93). The highest genetic distance between the parental lines was 0.56 and the lowest was 0.05, while the mean was 0.47. The highest genetic distance (0.56) was found between parental lines CKDHL0089, CML443 and CB323. The lowest genetic distance (0.05) was found between inbred lines I-42, and I-40. The majority (92%) of the genetic distance values fell between 0.40 and 0.60, suggesting the genotypes were moderately and distantly related (Figure 3). Cluster I consisted of 10 parental lines and further sub-divided into two sub-clusters. Cluster II also had two sub-clusters comprising of 15 parental lines, while cluster III consisted of five parental lines. Overall, the cluster analysis was effective in discriminating the parental lines into groups and in providing genetic information for breeding and conservation. In this analysis, three sets of parental lines with different genetic backgrounds were included.

**Figure 2:**
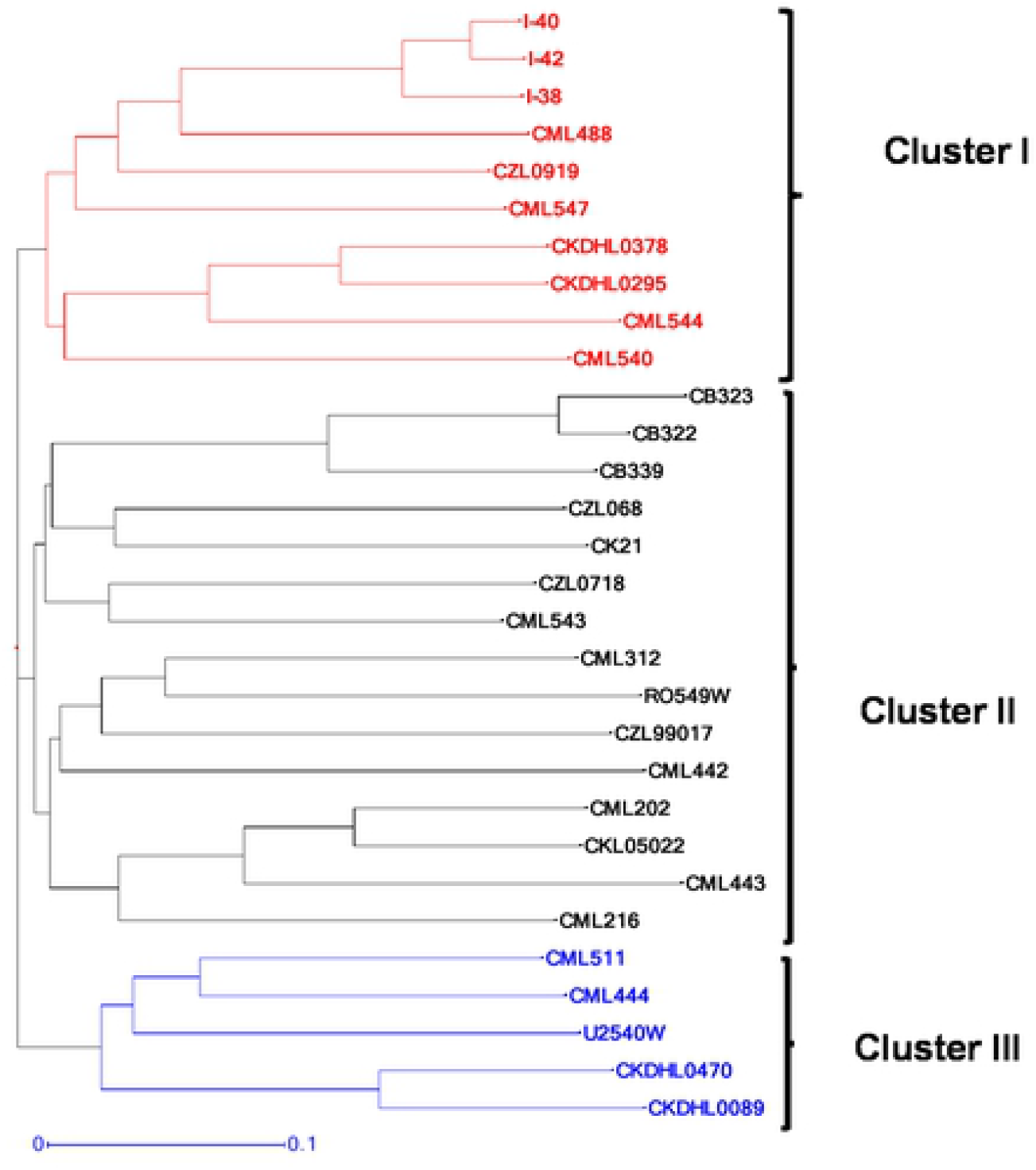
Neighbor-joining dendrograms based on UPGMA genetic dissimilarity depicting the genetic relationship between 30 parental maize lines based on 92 SNPs

**Figure 3:**
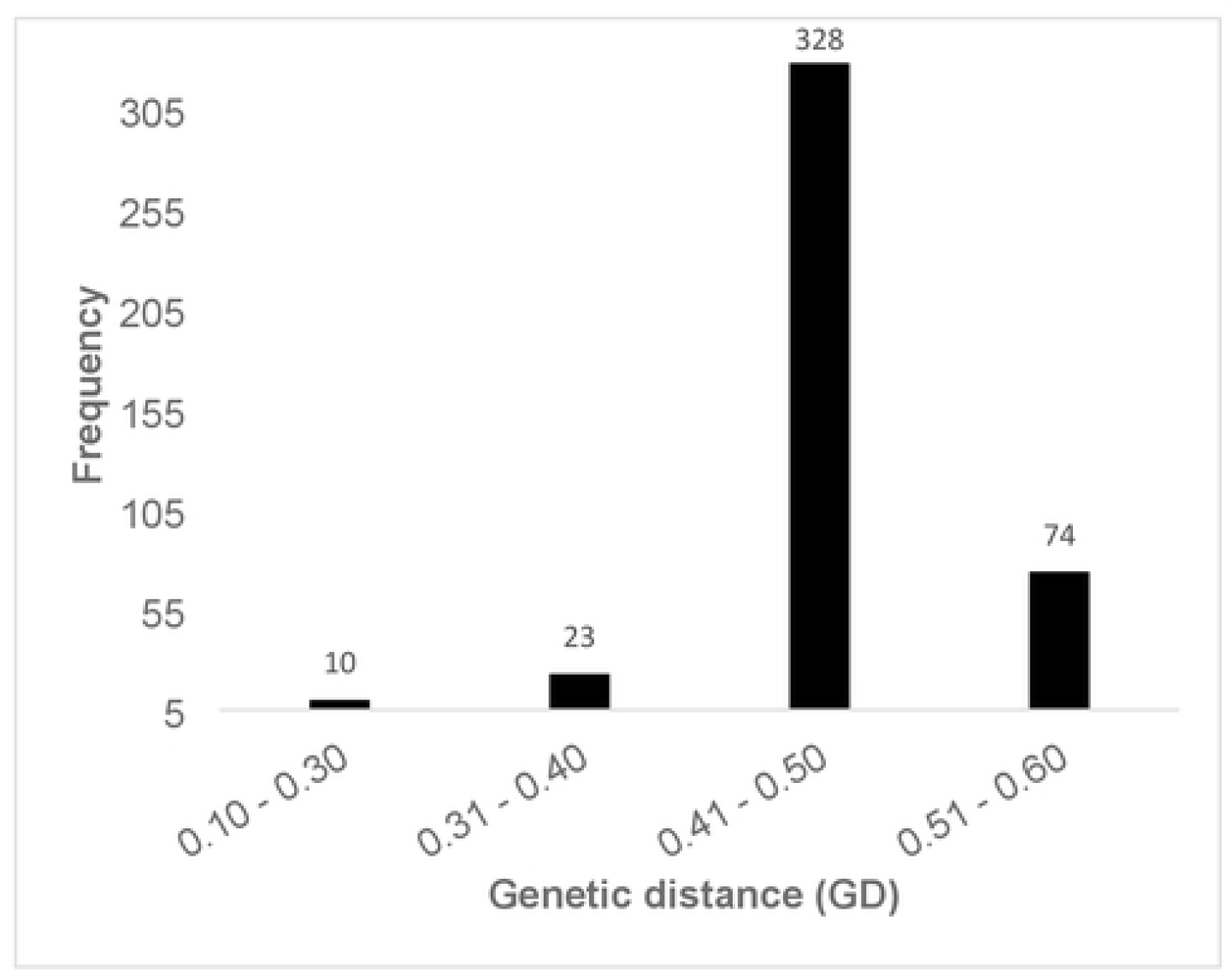
Genetic distance data summary of 30 parental maize lines using 92 SNPs

The clustering patterns of the parental lines were according to their genetic background. All the CB inbred lines (CB323, CB322 and CB 339) were grouped in Cluster II sub-cluster 1, while all the I inbred lines (I-40, I-42, I-38) were clustered in Cluster I sub-cluster 1. Similarly, all the CZL lines (CZL068, CZL0718 and CZL99017) except CZL0919 were clustered in Cluster I. The two CML heterotic tester lines, CML444 and CML202, which belong to heterotic groups A and B, respectively, were assigned in different clusters.

### 1.3.4 Parent-offspring test and grain yield performance for the selected maize hybrids

Parent offspring verification test revealed that out of 158 single-cross hybrids tested,16 hybrids had 0% contamination, 96 hybrids registered contamination level within the range of 0.54% to 4.89% (<5%) and 46 hybrids had contamination greater than 5% (Supplementary Table 3). Further quality analysis revealed that some of the ten contaminated lines were used for 90 single-cross hybrids. The test also confirmed that the remaining 68 single-cross hybrids were generated using pure parental inbred lines with acceptable genetic contamination (<5%).

The hybrids were evaluated for grain yield performance and the top and bottom 10% performing hybrids are presented in Table 3. Grain yield observed from 158 single-cross hybrids ranged from 2.83 t ha^-1^ for SCHP115 to 7.33 t ha^-1^ for SCHP17 and a mean yield of 5.55 t ha^-1^ was observed. The parent-offspring test was performed based on the criteria of < 5% genetic contamination on at least one of the parents and their hybrids. Based on the above criteria, of the 15 top performing hybrids, seven hybrids (SCHP29, SCHP95, SCHP94, SCHP134, SCHP44, SCHP114 and SCHP126) passed the test and represented 47% of the hybrids. Similarly, among the bottom 15 performing hybrids, five hybrids (SCHP113, SCHP102, SCHP32, SCHP71, and SCHP15) fulfilled the requirement. Notably, hybrids SCHP29 and SCHP115 among the top 10% and bottom 10% performing hybrids, respectively, exhibited genetic contamination of 0%.

**Table 3:**
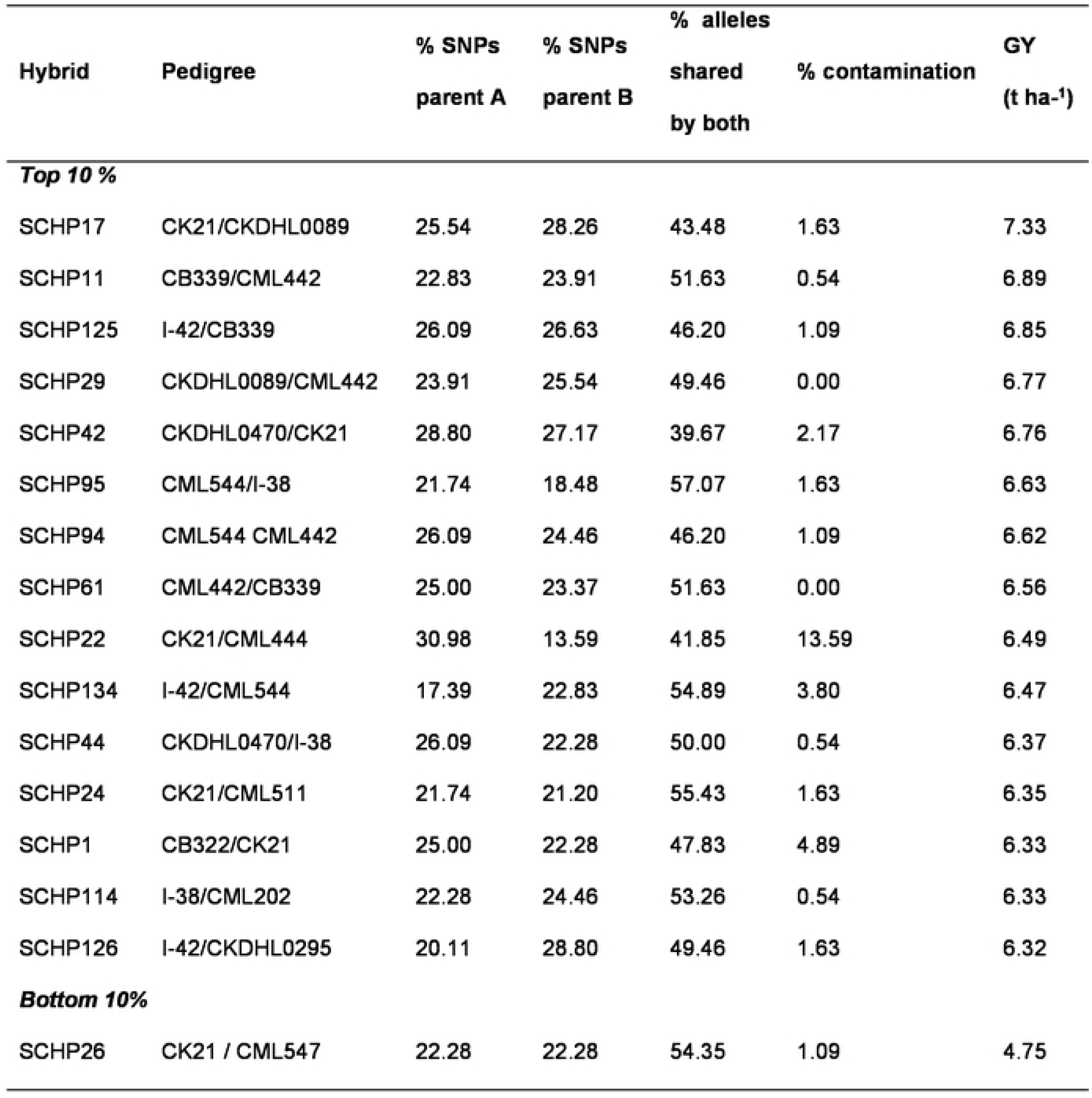
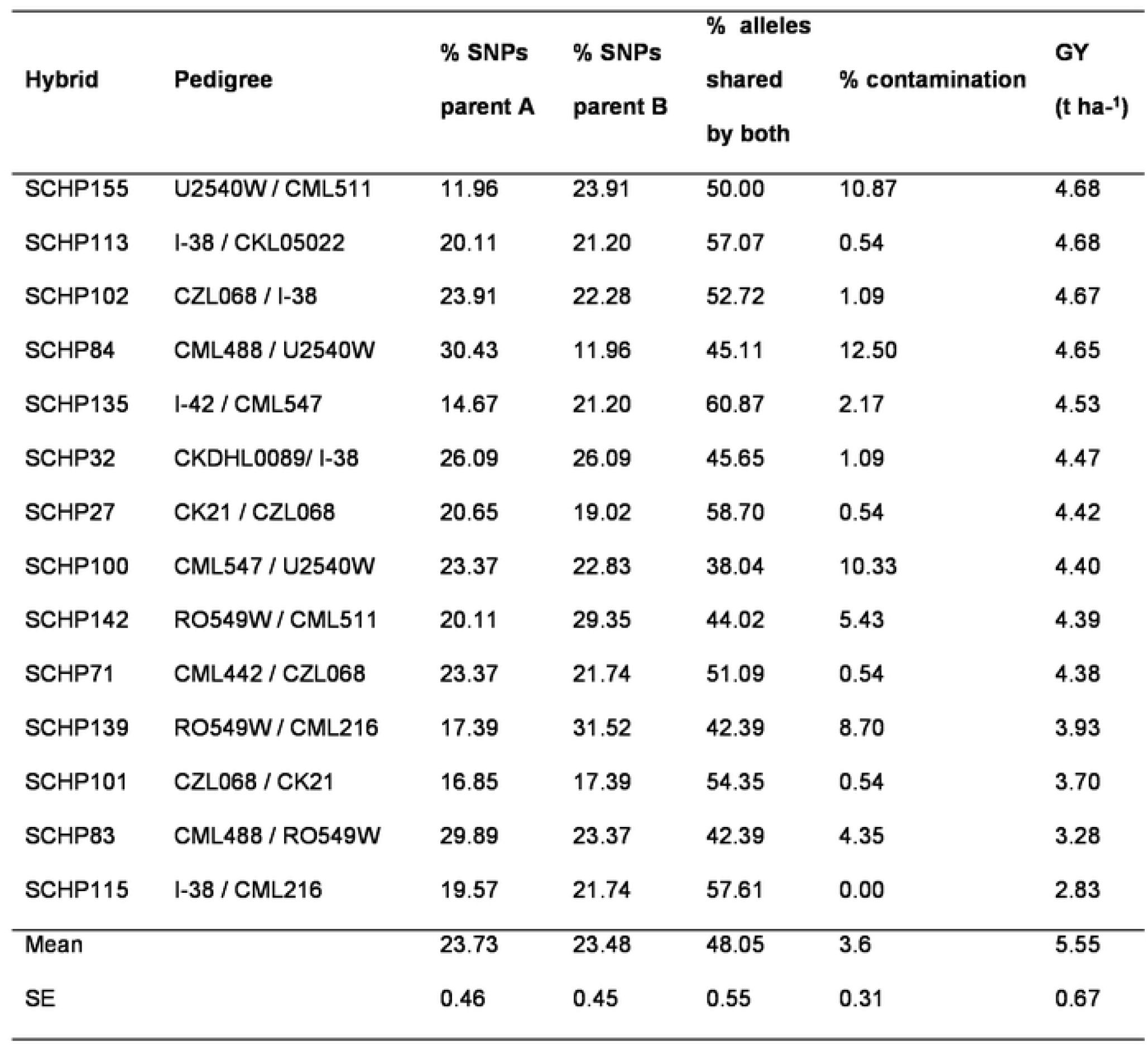
Parent-offspring test for top and bottom 10% performing hybrids based on grain yield

## 1.4 Discussion

In cross-pollinated crops such as maize, maintaining genetic purity for the inbred lines and the hybrids is vital for successful hybrid breeding and seed production. Significant changes in the genetic constitution of the inbred lines affect both the quality of newly developed hybrids and hybrid seed. The routine quality control genotyping in this study was done to detect any contamination which could have happened during hybrid development and inbred lines maintenance. According to Semagn et al. [2], inbred lines are regarded pure or fixed if the proportion of heterozygous SNP loci does not go beyond 5%. Inbred lines are also expected to maintain all the genetic characters that the breeder selected them for. Majority (67%) of the tested inbred lines were pure and fixed and 18 of them attained 100% genetic purity including four DH lines. The fact that all the DH lines (CKDHL0089, CKDHL0295, CKDHL0378 and CKDHL0470) used in this study exhibited 100% homozygosity indicates the advantages of using the DH approach in inbred line development. The DH approach enhances breeding efficiency through rapid generation of homozygous lines which are more fixed and predictive than those developed using conventional methods [11, 12]. This approach significantly shortens the breeding cycle though rapid development of fixed lines in two to three generations unlike the conventional approach that requires six to eight generations of inbreeding with approximately 99% homozygosity.

The remaining 14 inbred lines (CB323, CML202, CML216, CML442, CML443, CML511, CML544, CZL068, I-38, 1-42, CML 540, CZL99017, CML312 and I-40) with 100% genetic purity were generated through conventional system. This affirms that the maintenance of these inbred lines was carefully done for several generations of selfing for seed increase and they are suitable for high quality hybrid production. Similar findings were reported by Dao [13] in his study using 1237 SNPs, where the majority of the inbred lines tested exhibited 100% homozygosity. However, Ertiro et al. [14] reported that of the 265 inbred lines tested using 22,787 SNPs, only 22% of the inbred lines had 99.9% genetic purity. The higher levels of genetic purity observed in majority of the inbred lines used in this study indicates that the ARC-GCI’s maize breeding program is efficient and quality oriented in terms of inbred line development and maintenance. However, inbred line CK21, which is currently used as a parent in some of the experimental hybrids had residual heterozygosity of 5.43%, which is slightly higher than the threshold of 5%. It should, therefore, be purified using ear-to-row selection methods. Lines with more than 15% residual heterozygosity are likely to have been contaminated with pollen from unrelated genetic materials and should be discarded [3]. The higher level of heterogeneity observed in some of the inbred lines may be attributed to either that the inbred line is in the early generation of inbreeding or there was pollen contamination and/or seed admixture during maintenance breeding. An additional generation of inbreeding and extensive selection is highly recommended in order to fix these inbred lines. Some reports suggest that due to the strong inbreeding depression, higher levels of residual heterozygosity may have been deliberately maintained at early breeding level. Ertiro et al. [14] also reported higher levels of residual heterozygosity among the inbred lines tested from Ethiopian Institute of Agricultural Research (EIAR) due to use of early generation inbred lines (S4). Warburton et al. [15], further suggested that high level of residual heterozygosity may occur due to pollen contamination and/or seed mixture during seed regeneration, maintenance and bulking.

The rationale for doing parent-offspring test is to confirm if the particular hybrid is the true resultant F1 hybrid derived from the original inbred lines with no pollen contamination or within acceptable contamination levels [3]. Thus, the test provides a means to check if the pollination was done correctly during hybrid development. During hybridisation, there is a possibility of pollen contamination arising from self-pollination or cross-pollination from undesired neighbouring crops due to inadequate isolation distance. Therefore, parent-offspring test is important to ensure production of genuine quality hybrid seed.

Out of 158 experimental hybrids tested, 90 failed the test due to higher percentage of contamination (greater than 5%). This could be attributed partly to the use of genetically impure parental inbred lines or partly due to pollen contamination during hybrid production. The results revealed that the resultant hybrids derived from segregating inbred lines exhibited higher levels of genetic contamination of greater than 5%. Daniel et al. [16] reported a similar trend, where inbred lines with higher percentage of residual heterozygosity resulted in hybrids with higher contamination percentage. The results clearly confirm that the purity level of the parental inbred lines determines the purity of the resultant hybrids. In addition, it was evident that there was genetic contamination due to lack of pollen control between the crossing block and the neighbouring field as hybrids developed from pure inbred lines also revealed high levels of contamination. This could be due to failure in observing adequate isolation distance of not less than 360 m recommended in hybrid seed production. Genetic contamination of the experimental hybrids could also be as a result of incomplete detaselling as was postulated by MoreiraI et al. [17]. However, this kind of impurity is usually tolerated at a level of 3-5%, without any effect on the yield performance. Thus, the low level of hybrid performance detected in this study could be due to the use of the genetically impure inbred parental lines. The finding concurs with Ipsilandis et al. [18], who reported a significant influence of seed purity status to the yield performance of different genetic materials. This further confirms that pure hybrid seed has better competitive ability and yields better than low purity seed.

Polymorphic information content (PIC) gives an estimate of the discriminatory ability and effectiveness of molecular markers with respect to the number of alleles that are expressed and their relative frequencies [19]. Lander and Botstein [20] described PIC mean value of >0.50 as highly informative, 0.25-0.50 moderately informative and <0.25 is slightly informative. Hence, the mean PIC value of 0.44 observed in this study for the inbred lines confirm that the markers used were effective in discriminating the genotypes, reasonably informative and of good quality. This value is on the higher side compared to PIC values reported in some of the past related studies [21,22]. Hao et al. [23] reported PIC values within the range of 0.01 to 0.38 using 1536 SNP markers on 95 parental inbred lines in maize. In another study, Yang et al. [24] reported PIC values ranging from 0.27 to 0.38 with a mean of 0.34 while using 884 SNP markers. The mean PIC value observed in this study is comparable to the one reported by Adeyemo and Omidiji [25] of 0.43 using 66 SSR markers. The disparity between the mean PIC observed in this study and the findings of earlier scientist could be attributed to differences in the composition of the genetic materials, sample size as well as the choice of number of markers used in the studies. The good quality of the 92 SNPs used in this study could be attributed to the fact that they were a subset of the 100 SNPs distributed across the 10 chromosomes in the maize genome, which were carefully selected and recommended by CIMMYT for quality control genotyping.

The mean observed heterozygosity (H_0_) of 0.09 and mean inbreeding coefficient of 0.80 reported in this study among the inbred lines indicate that the majority of the lines attained an appreciable level of homozygosity and were fixed. The low level of residual heterozygosity and high inbreeding coefficient in the present study further confirmed the existence of high level of inbreeding [26], which was expected among the parental inbred lines. The hybrids on the other hand, revealed very low F_IS_ values ranging from −0.02 to 0.17, with a mean value of −0.05 validating an excess of heterozygotes, which was expected for hybrids.

It is critical that genetic relatedness of parental inbred lines be established to facilitate efficient exploitation of heterosis in hybrid breeding. The use of distanced based cluster approach to partition parental lines into distinct clusters is a reliable and appropriate approach to measure relatedness among individuals within a population [27]. Higher genetic distance of 0.56 observed between any combination of the following three parental inbred lines namely, CKDHL0089, CML443 and CB323 indicate that these lines originate from different ancestors while lowest genetic distance of 0.05 between lines I-42 and I-40 suggest that these are sister lines with shared pedigree across most generations. On average, majority of the inbred lines studied (92%) had their genetic distance values falling within the range of 0.4 to 0.6. In a related study, Boakyewaa Adu et al. [28] reported that on average, 67.7% of 94 maize inbred lines had genetic distances within the range of 0.32 to 0.42. CIMMYT conducted a similar study on 450 inbred lines and 95% of the pair of inbred lines showed genetic distances ranging between 0.3 and 0.5 [2]. Genetic distances among the studied inbred lines revealed the uniqueness of the studied lines and presence of substantial genetic variability that could be exploited in the breeding program.

## 1.5 Conclusion

This study showed that the set of SNP markers recommended for quality control test were effective and reliable in assessing genetic purity. The results of study will be helpful in the verification of genetic purity of maize hybrid seed developed by ARC-GCI. The inbred lines used in the present study were expected to be genetically pure with not more than 5% residual heterozygosity, but 33% of inbred lines showed residual heterozygosity of greater than 5% which requires additional generations of purification. Parent-offspring test conducted on 158 experimental hybrids led to the elimination of 60% hybrids since at least one of their parental inbred lines failed the genetic purity test. Of the 30% of the hybrids that passed the quality control test, seven high yield potential hybrids were recommended for further evaluation and release. Failure of some genotypes to pass inbred line genetic purity test and parent-offspring test suggests the need for further quality improvement by ARC-GCI maize-breeding program in the breeding nurseries and during pollinations for hybrid production.

## 1.6 Acknowledgements

The authors would like to thank the Alliance for a Green Revolution in Africa (AGRA) for providing funding for this study (grant number 2014PASS013), ARC-GCI maize-breeding program for providing the experimental materials and an internship opportunity to the first author.

## 1.7 Conflicts of Interest

The authors declare that they have no known competing interests that could influence the work reported in this paper.

## 1.8 Author Contributions

Conceptualization: Chimwemwe Josia, Julia Sibiya and Kingstone Mashingaidze

Data curation: Chimwemwe Josia, Assefa B. Amelework, Cousin Musvosvi and Aleck Kondwakwenda

Formal analysis: Chimwemwe Josia, Assefa B. Amelework, Cousin Musvosvi and Aleck Kondwakwenda

Funding acquisition: Julia Sibiya and Kingstone Mashingaidze

Project administration: Julia Sibiya and Kingstone Mashingaidze

Resources: Julia Sibiya and Kingstone Mashingaidze

Supervision: Julia Sibiya and Kingstone Mashingaidze

Validation: Julia Sibiya, Kingstone Mashingaidze and Assefa B. Amelework

Writing – original draft: Chimwemwe Josia, Aleck Kondwakwenda and Cousin Musvosvi

Writing – review & editing: Chimwemwe Josia, Aleck Kondwakwenda, Julia Sibiya, Kingstone Mashingaidze, Assefa B. Amelework and Cousin Musvosvi

